# From simplicity to complexity: The gain or loss of spot rows underlies the morphological diversity of three *Drosophila* species

**DOI:** 10.1101/2020.04.03.024778

**Authors:** William A. Dion, Mujeeb O. Shittu, Tessa E. Steenwinkel, Komal K. B. Raja, Prajakta P. Kokate, Thomas Werner

## Abstract

To understand how novel animal patterning emerged, one needs to ask how the development of color patterns has changed among diverging species. Here we examine three species of fruit flies – *Drosophila guttifera* (*D. guttifera*), *Drosophila palustris* (*D. palustris*), and *Drosophila subpalustris* (*D. subpalustris*) – displaying a varying number of abdominal spot rows that were either gained or lost throughout evolutionary time. Through *in situ* hybridization, we examine the mRNA expression patterns for the pigmentation genes *Dopa decarboxylase* (*Ddc*), *tan* (*t*), and *yellow* (*y*) during pupal development. Our results show that *Ddc*, *t*, and *y* are co-expressed in identical patterns, each foreshadowing the adult abdominal spots in *D. guttifera*, *D. palustris*, and *D. subpalustris*.

## 1. Introduction

The complexity and diversity of animal body coloration in the natural world are astounding. Unique patterns like cheetah spots and zebra stripes beg the question – how did these traits evolve? To understand how novel morphologies arose, one needs to ask how alterations to organismal development occurred over evolutionary time (Raff, 2000). Butterfly wings have served as a system to unravel the molecular mechanisms underlying complex pattern development (Carroll et al., 1994; Matsuoka and Monteiro, 2018; Monteiro et al., 2013; Zhang and Reed, 2016; Zhang et al., 2017), and the examination of American cockroaches, large milkweed bugs, and twin-spotted assassin bugs progressed the knowledge of the process of body coloration (Lemonds et al., 2016; Liu et al., 2014; Zhang et al., 2019). Moreover, pigmentation has been shown to be vital to the lifecycles of agricultural pests and human disease vectors, such as the Asian tiger mosquito, black cutworm, brown planthopper, and kissing bug (Chen et al., 2018; Lu et al., 2019; Sterkel et al., 2019). However, these studies were built upon the robust knowledge of pattern and pigmentation development gained through the study of fruit flies, in particular, *Drosophila melanogaster* (*D. melanogaster*).

Our initial understanding of the genetic mechanisms responsible for the process of adult fruit fly pigmentation began with the study of *D. melanogaster* (Wright, 1987; Karlson and Sekeris, 1962). Very recent studies have examined the relationship between pigmentation and thermal plasticity (De Castro et al., 2018; Gibert et al., 2017), and how pigmentation of the sex comb contributes to *Drosophila* mating success (Massey et al., 2019). Investigating how pigmentation develops in *D. melanogaster* provided the foundation to understand the same processes in other fruit flies. This knowledge, in turn, has facilitated studies of species divergence (Lamb et al., 2020) and positioned *Drosophila* pigmentation as a model to study how gene-regulatory networks – the regulatory mechanisms responsible for organismal development (Davidson and Levin, 2005) – evolved (Camino et al., 2015; Gibert et al., 2018; Grover et al., 2018; Ordway et al., 2014; Rebeiz and Williams, 2017; Roeske et al., 2018). The *Drosophila* pigmentation pathway with the enzymes and reactions necessary to produce black, brown, and yellow coloration seen on the bodies of fruit flies, is shown in Figure 1 (Rebeiz and Williams, 2017; True et al., 2005; Wittkopp et al., 2003).

**Figure 1:**
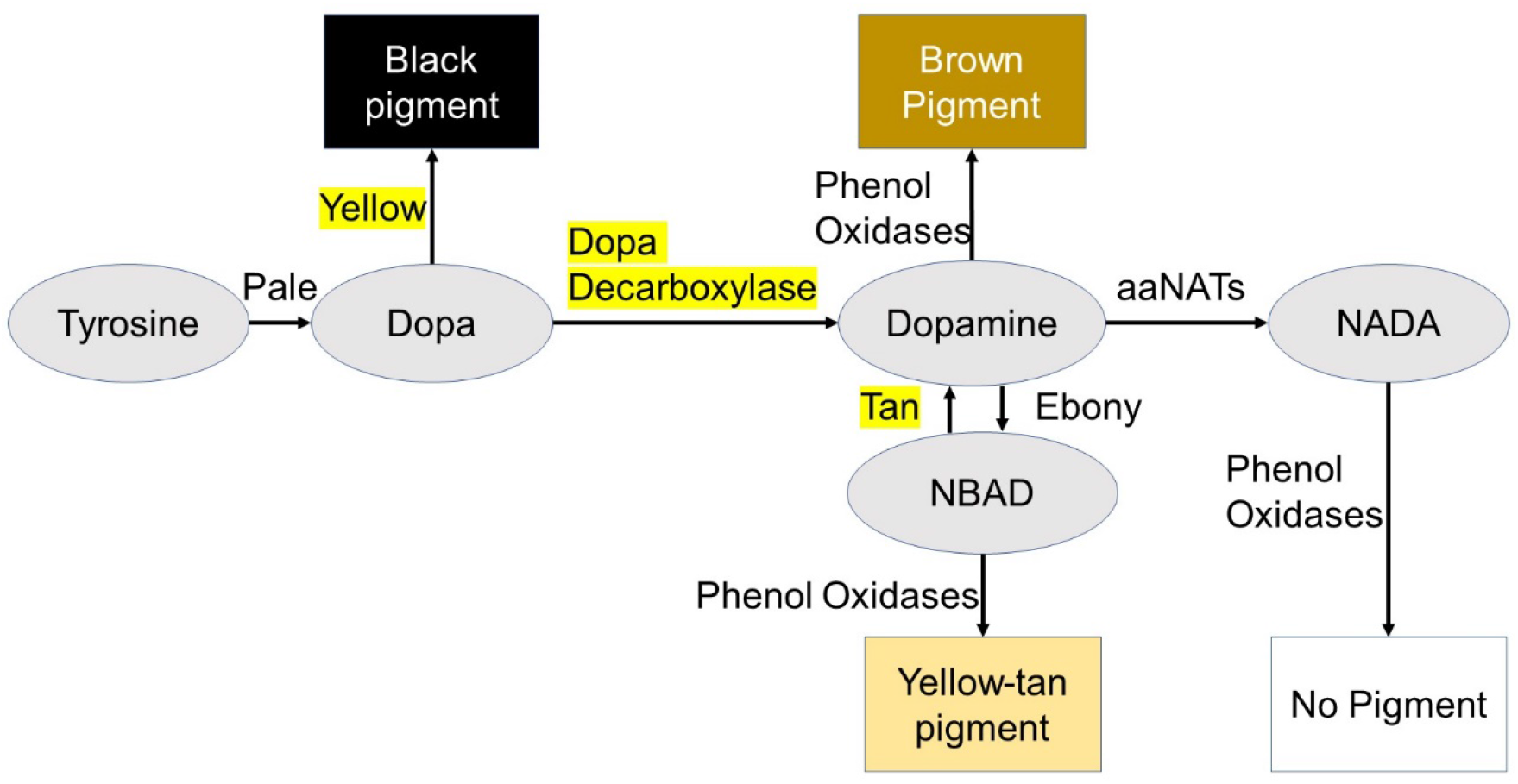
The pigmentation pathway of *Drosophila*. This illustration of the pigmentation pathway is adopted from (Rebeiz and Williams, 2017; True et al., 2005; Wittkopp et al., 2003). Tyrosine is converted to dopa by Pale. Dopa can be shunted into black pigment by Yellow (encoded by *y*) or further converted into dopamine by Dopa decarboxylase (encoded by *Ddc*). Dopamine proceeds one of three ways: it can become brown pigment through the activity of phenol oxidases; it can be converted into *N–*acetyl dopamine (NADA) through arylalkylamine *N-*acetyl transferases (aaNATs) and thus result in a lack of pigmentation through the activity of phenol oxidases; or it may become *N-*β-alanyl dopamine (NBAD) through the activity of Ebony, followed by a transition to a yellow-tan pigment by phenol oxidases. The protein Tan (encoded by *t*) functions opposite of Ebony by converting NBAD into dopamine, which may result in brown pigment. The gene products for *Ddc*, *t*, and *y* are highlighted.

While the process of *Drosophila* pigmentation patterning involves many genes, our study focuses on three: *Ddc*, *t*, and *y*, which are all essential for the production of black and brown coloration. *Ddc* is integral to the development of *Drosophila* pigmentation, with the mutant phenotype lacking the dark coloration seen on the wild type fly (Walter et al., 1996; Wright et al., 1976). The genes *t* and *y* are also required for melanization. Mutants of the *t* gene exhibit a tan as opposed to a black body pigmentation, while *y* mutants display a yellow body color (Biessmann, 1985; Hotta and Benzer, 1969; Kornezos and Chia, 1992; True et al., 2005).

*D. melanogaster* has a relatively simple abdominal pigmentation pattern, as compared to other *Drosophila* species. The quinaria group, an adaptive radiation of non-model fruit flies, displays a great variety of abdominal and wing pigmentation patterns (Bray et al., 2014; Werner et al., 2018). This abundant morphological diversity and the recent divergence of the lineage (20 million years ago (Scott Chialvo et al., 2019)) allows for the identification of molecular mechanisms underlying differences in species morphology. One member of the quinaria group, *D. guttifera*, is already becoming established as a model to study complex pattern development (Fukutomi et al., 2020; Koshikawa et al., 2015; Koshikawa et al., 2017; Werner et al., 2010).

The abdominal spot pattern of *D. guttifera* consists of six rows of spots: three rows on the left side (dorsal, median, and lateral row), which are mirrored on the right side of the abdomen. The pattern of *D. palustris* is similar but lacks the dorsal pair, while in *D. subpalustris*, only the lateral pair of spot rows is present (Figure 2). In addition to displaying spots, the abdomens of each of the three fruit fly species exhibit wide areas of dark shading. *D. guttifera* shows two somewhat distinct areas of shading: a wide swath encompassing the spotted region that is shared by all three species, plus a specific dorsal midline shade. Furthermore, *D. guttifera* shows blackish stripes along the dorsal segment boundaries, which are absent in the other species.

**Figure 2:**
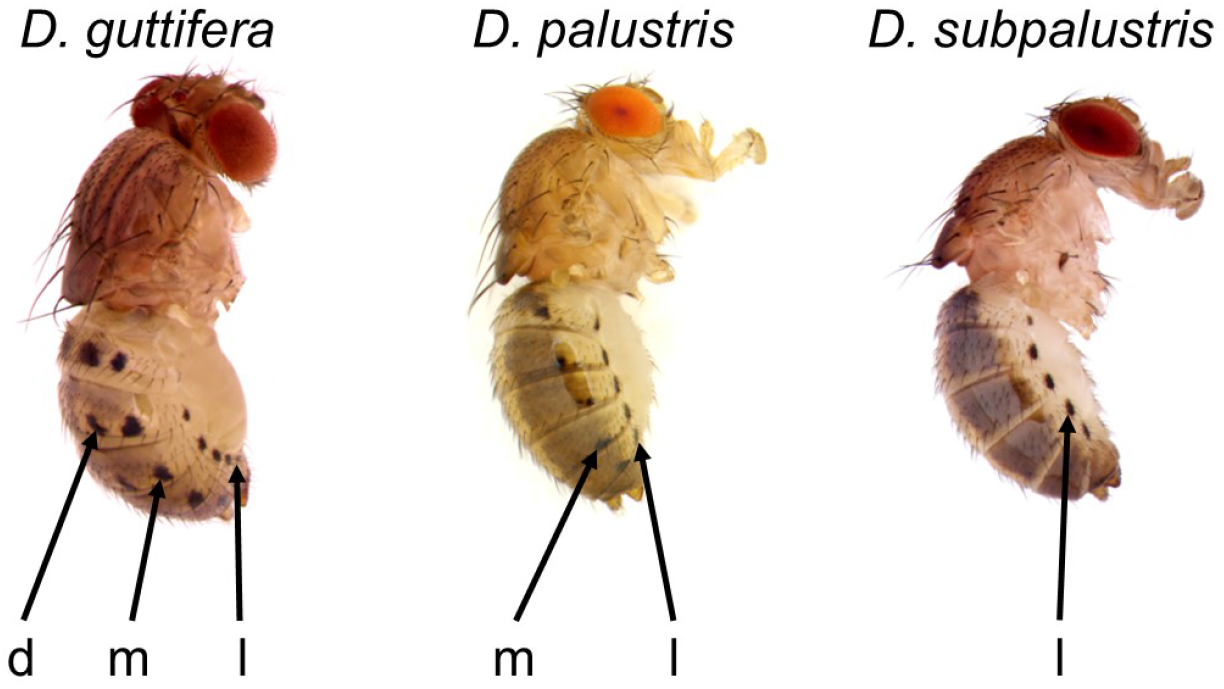
Spot pattern complexity in the quinaria group. Three members of the quinaria group are shown from a lateral view. The dorsal (d), median (m), and lateral (l) rows of spots are labeled. Images are from (Werner et al. 2018).

In the current study, we show that abdominal color pattern diversity among the quinaria group members *D. guttifera*, *D. palustris*, and *D. subpalustris* is strictly modular and that *Ddc*, *t*, and *y* are co-expressed in identical patterns where dark spots will appear.

## 2. Materials and Methods

### 2.1 *Drosophila* stocks –*D. guttifera*, *D. palustris*, and *D. subpalustris*

*D. guttifera* and *D. subpalustris* were purchased from the *Drosophila* Species Stock Center, stock numbers 15130 – 1971.10 and 15130 – 2071.00, respectively. We collected *D. palustris* in Waunakee, Wisconsin. All fly stocks were maintained at room temperature on cornmeal-sucrose-yeast medium (Werner et al., 2018).

### 2.2 Identification of pupal stages

Pupal developmental stages were determined according to (Bainbridge and Bownes, 1981) and (Fukutomi et al., 2017).

### 2.3 *in situ* hybridization probe design for *Ddc*, *t*, and *y*

RNA *in situ* hybridization probes were 200 to 500 bases in length. The *Ddc* probe for *D. guttifera* and the *t* probe used for all three species were synthesized from a *D. guttifera*-derived DNA template, while the *Ddc* and *y* probes for both *D. palustris* and *D. subpalustris* were transcribed from *D. palustris* DNA and used interchangeably. We used Mean Green PCR Master Mix (Syzygy Biotech Solutions) to amplify the partial coding regions with forward and reverse primers (Table 1). The PCR products were extracted and purified with a Thermo Scientific GeneJET Gel Extraction Kit and cloned into the pGEM-TEasy vector, using *E. coli* DH5α cells. Colony PCR with the M13 forward and reverse universal primer pair was used for screening, and the Thermo Scientific GeneJET Plasmid Miniprep Kit was used for plasmid purification. The insertion direction into the pGEM-TEasy vector was determined through PCR with the M13 forward universal primer and either the internal forward or internal reverse primer (Table 1). Depending on the insertion direction, either SP6 or T7 RNA polymerase was used to produce a DIG (digoxigenin)-labeled RNA anti-sense probe (Roche DIG RNA Labelling Kit (SP6/T7)). GenePalette was used for computational biology (Rebeiz and Posakony, 2004).

**Table 1:**
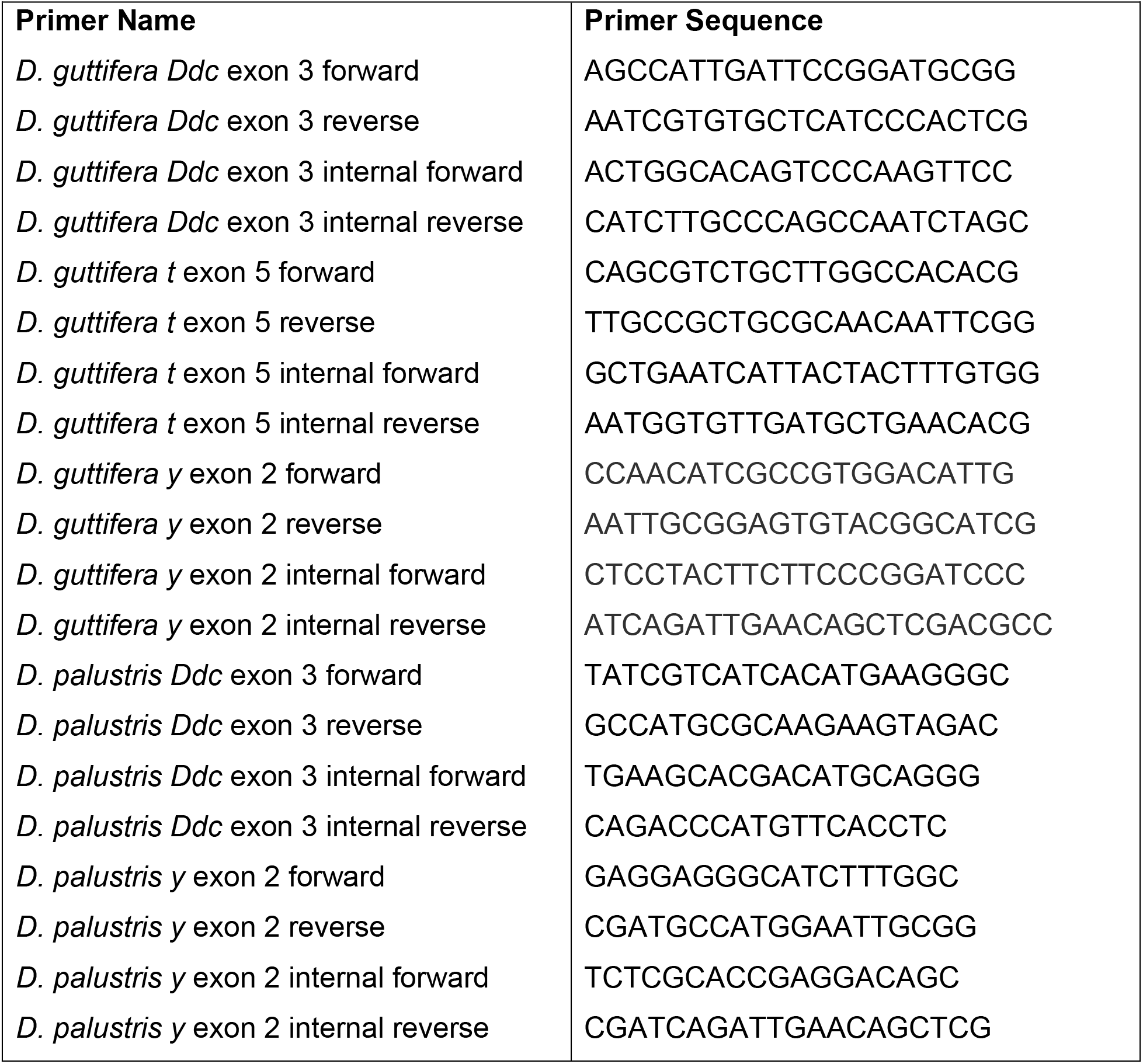
Primers.

### 2.4 Preparation of pupae for RNA *in situ* hybridization

When pupae had developed to the desired stage, they were cut along the anterior-posterior axis either between the eyes or on their side through the eyes. The pupal halves were fixed in 4% paraformaldehyde (Electron Microscopy Sciences) and kept at −20°C in pure ethanol.

### 2.5 *in situ* Hybridization of the pupae

The *in situ* hybridization procedure was adopted from (Jeong et al., 2008). The tissues were washed between each step with PBST. On the first day, pupae were treated with a 1:1 xylenes to ethanol mixture to remove residual fat tissue. The pupal tissue was then fixed (4% paraformaldehyde), treated with Proteinase K for 10 to 15 minutes (1:25,000 dilution of a 10 mg/mL stock solution), fixed again (4% paraformaldehyde), and then incubated with the anti-sense RNA probe (1:500 dilution) for 18 to 72 hours at 64°C to 65°C. Pupae were gently agitated periodically. The pupae were then incubated in Roche α-DIG AP Fab Fragments (1:6000) at 4°C overnight. On the final day, the tissues were incubated with the BCIP/NBT staining solution (Promega) in the dark until patterns were fully developed (approximately two to 18 hours).

### 2.6 Imaging of *Ddc*, *t*, and *y* expression patterns after *in situ* hybridization

z-Stacks of images were taken with Olympus cellSens software, using an Olympus SZX16 microscope and an Olympus DP72 camera. The digital images were stacked with Helicon Focus software.

## 3. Results

### 3.1 *D. guttifera* pattern development

The expression patterns of *Ddc*, *t*, and *y* during pupal development foreshadowed the abdominal spots of *D. guttifera*. Although the expression patterns were spatially the same for the spots, they differed temporally. *Ddc* mRNA was detected at pupal stages P10 through P13, *t* mRNA at P11 and P12, and *y* mRNA at P10 (Figure 3). For the rest of the pattern, only *y* expression correlated with both the dorsal midline shade and intersegment stripes at stage P10 (Figure 4). However, we were unable to show any gene expression foreshadowing the broader shading around the dorsal and median spot rows.

**Figure 3:**
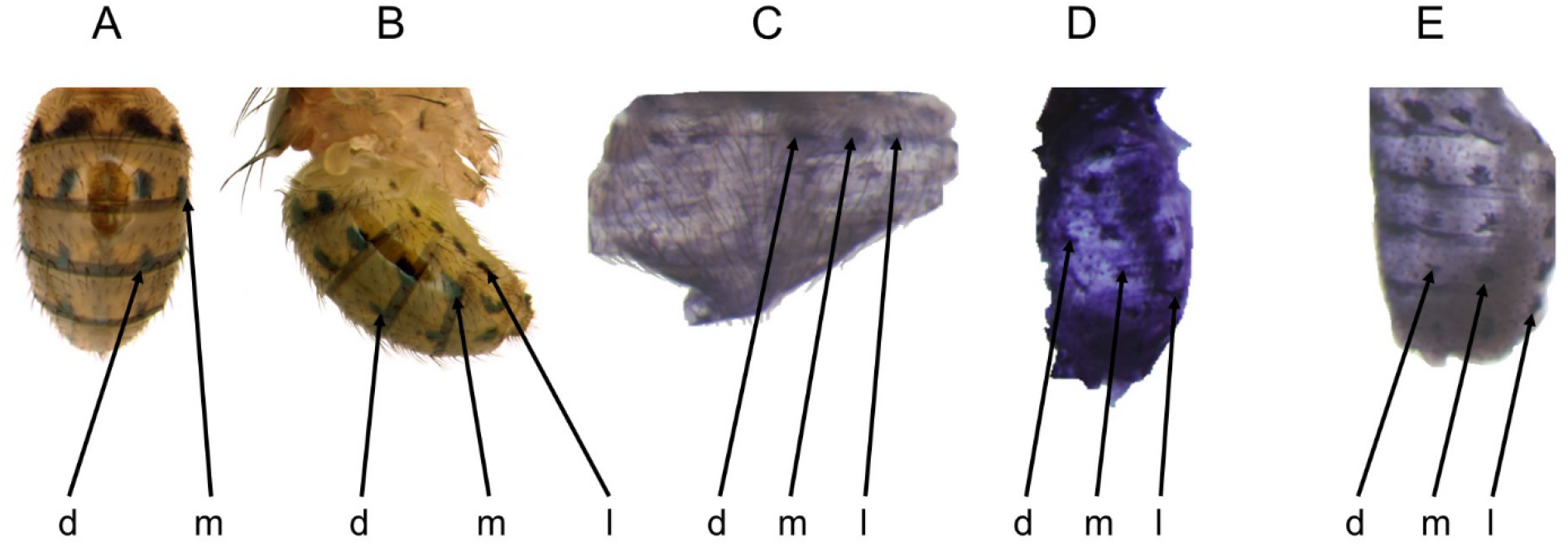
*in situ* Hybridization signals of *Ddc*, *t*, and *y* during *D. guttifera* pupal development foreshadowed the adult spot pattern. The spot rows are labeled as dorsal (d), median (m), and lateral (l). (A) Dorsal and (B) lateral view of adult *D. guttifera* (Werner et al., 2018). (C) *Ddc* mRNA expression at stage P13. (D) *t* mRNA at stage P11. (E) *y* mRNA expression at stage P10.

**Figure 4:**
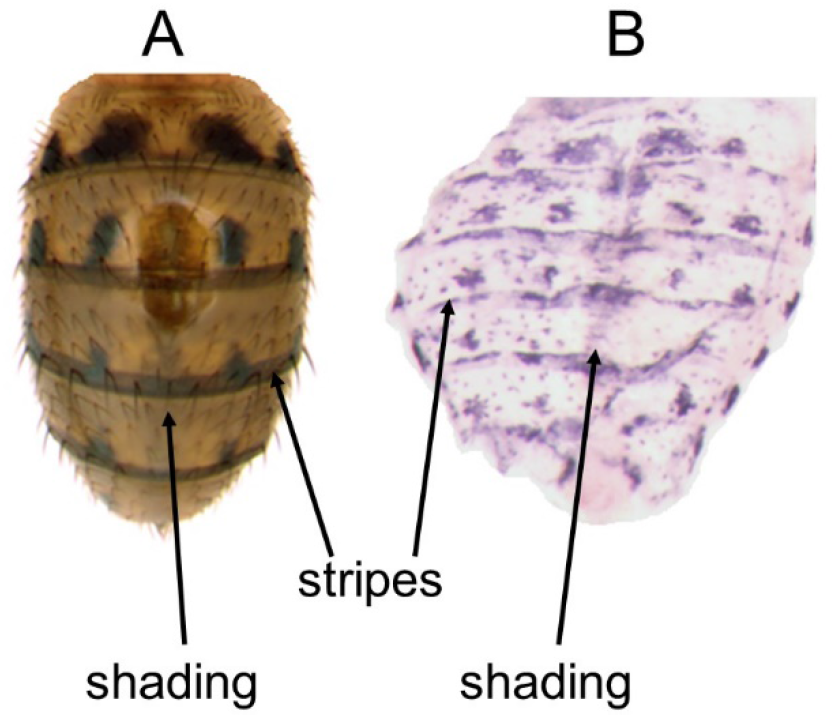
The *in situ* hybridization result of *y* during *D. guttifera* pupal development correlated with the adult abdominal dorsal midline shading and the intersegment stripes. (A) Dorsal view of adult *D. guttifera* (Werner et al., 2018). (B) *y* mRNA expression at stage P10 foreshadowing the dorsal midline shading (shading) and the intersegment stripes (stripes).

### 3.2 *D. palustris* pattern development

*D. palustris* lacks three components of the *D. guttifera* pattern: the dorsal rows of spots, the dorsal midline shade, and the intersegment stripes. Just as in *D. guttifera*, the mRNA expression patterns of *Ddc*, *t*, and *y* prefigured the *D. palustris* spot pigmentation. *Ddc* mRNA was present at stages P11 through P12, *t* at P12, and *y* at P10 and P12 (Figure 5). We observed a disparity in the mRNA expression patterns foreshadowing the median rows of spots, which correlated with the variation of median spot row pigmentation intensity seen on adult flies (Werner et al., 2018). However, only the expression of *t* mRNA at stage P12 correlated with the shading pattern (Figure 6).

**Figure 5:**
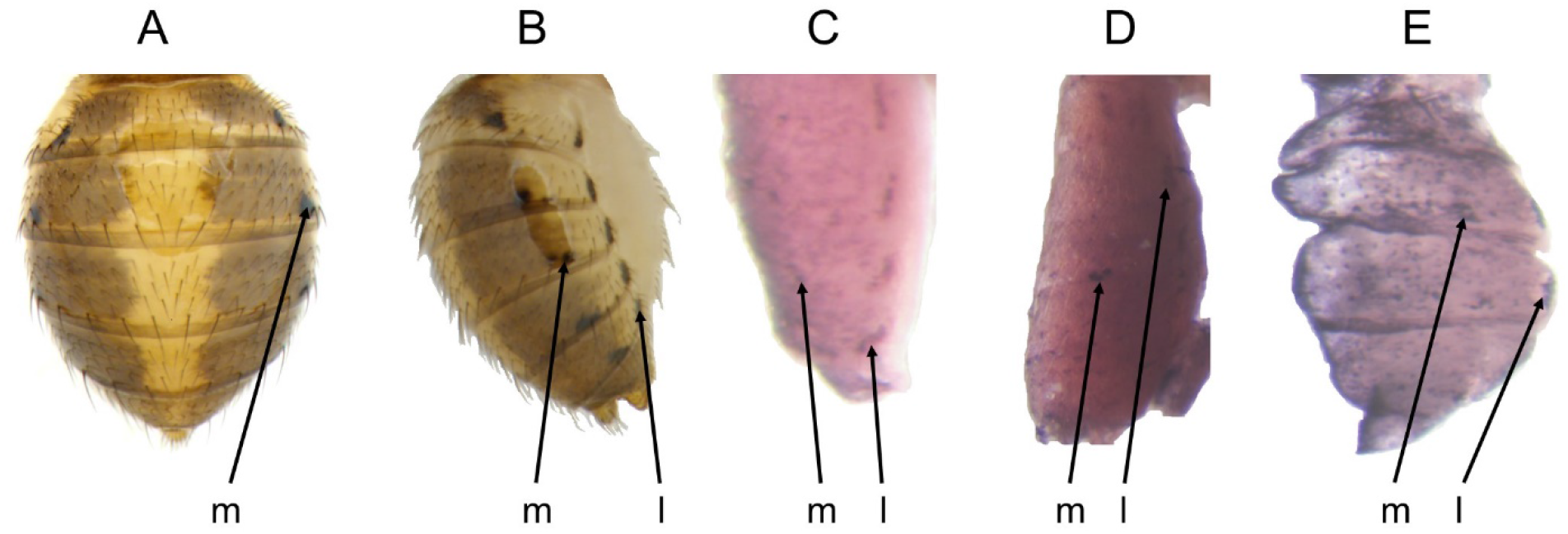
*in situ* Hybridization signals of *Ddc*, *t*, and *y* during *D. palustris* pupal development foreshadowed the abdominal spot pattern. The spot rows are labeled as median (m) and lateral (l). (A) Dorsal and (B) lateral view of adult *D. palustris* (Werner et al., 2018). (C) *Ddc* mRNA expression at stage P11. (D) *t* gene expression foreshadowing spots at a stage between P11 and P12. (E) *y* mRNA expression at stage P10.

**Figure 6:**
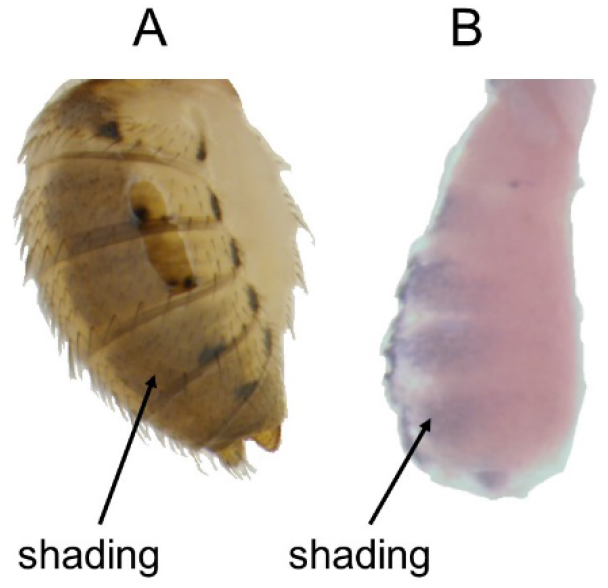
The *in situ* hybridization result of *t* during *D. palustris* pupal development correlated with the adult abdominal shading. (A) Lateral view of adult *D. palustris* (Werner et al., 2018). (B) *t* mRNA expression at stage P12.

### 3.3 *D. subpalustris* pattern development

*D. subpalustris* exhibits the simplest pattern: one pair of lateral spot rows and shading similar to that of *D. palustris*. The *Ddc*, *t*, and *y* expression patterns during pupal development foreshadowed the abdominal spots of *D. subpalustris*; *in situ* hybridization signals were seen for *Ddc* at stages P11 and P12, *t* at stage P12, and *y* at stage P10 (Figure 7). The shading pattern is prefigured by *Ddc* mRNA at stage P11 (Figure 8).

**Figure 7:**
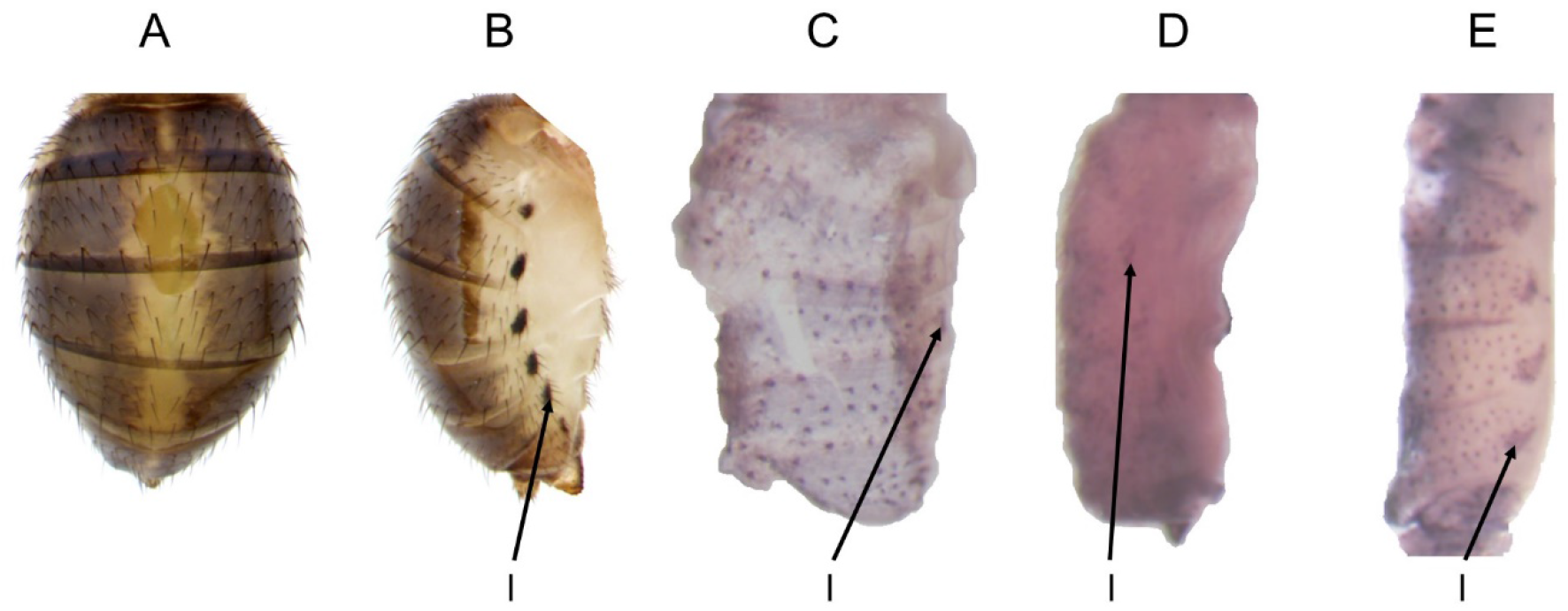
*in situ* Hybridization signals for *Ddc*, *t*, and *y* during *D. subpalustris* pupal development prefigured the abdominal spot pattern. The spot rows are labeled as lateral (l). (A) Dorsal and (B) lateral view of adult *D. subpalustris* (Werner et al., 2018). (C) *Ddc* gene expression foreshadowing spots at stage P11. (D) *t* gene expression at stage P12. (E) *y* mRNA expression at stage P10.

**Figure 8:**
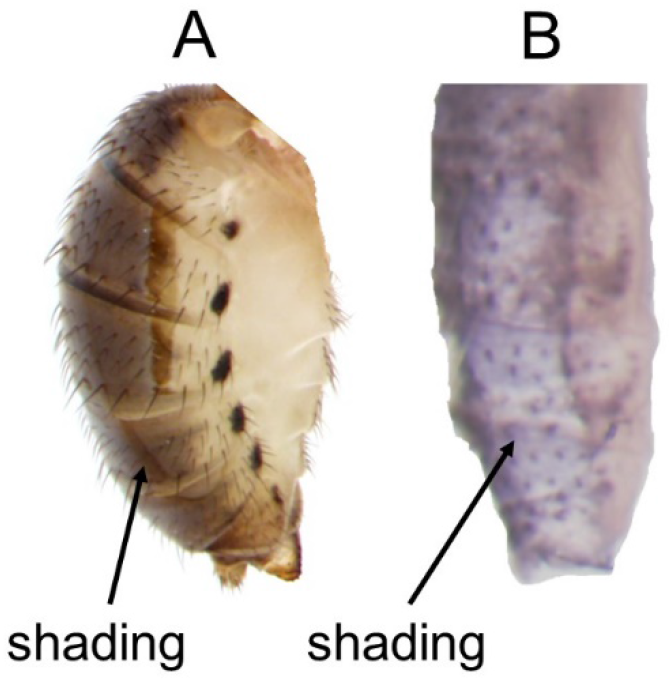
The *in situ* hybridization result for *Ddc* during *D. subpalustris* pupal development foreshadowed the adult abdominal shading. (A) Lateral view of adult *D. subpalustris* (Werner et al., 2018). (B) *Ddc* mRNA expression at stage P11.

## 4. Discussion

Here we show the developmental mechanisms underlying the complex pigmentation of three *Drosophila* species. *Ddc*, *t*, and *y* are spatially co-expressed in the developing abdomens, precisely foreshadowing the diverse dark spot patterns in three quinaria group species. Furthermore, the shades and intersegment stripes are uniquely foreshadowed by only one of the three genes: *Ddc* in *D. subpalustris*, *t* in *D. palustris*, and *y* in *D. guttifera*. These data indicate that over the past 20 million years since the divergence of this species group, the evolution of the same or similar gene-regulatory network(s) accounts for the difference in spot patterns, and that the shade patterns have evolved independently.

We demonstrate the potential of these three species as a model to understand color pattern diversity. The polka-dotted patterns are assembled from modules: a dorsal, median, and lateral pair of spot rows, as well as a wide shade on the left and right side of the abdomen (encompassing the median and dorsal spots), a dorsal midline shade, and intersegment stripes. Each species has either the complete set of modules (*D. guttifera*) or a subset of it (*D. palustris* and *D. subpalustris*).

The spot pattern diversity seen among the three non-model species alone position them as an emerging system to study color pattern diversity. The co-expression of multiple genes in complex spatial patterns is a phenomenon rarely reported in the literature; however, we show this event as responsible for the spot patterning of these three quinaria group species. Furthermore, each pair of spot rows behaves like a set of independent, serial homologs, similar to the repetitive pattern elements within butterfly wing sections (Monteiro 2008). The underlying gene co-expression and serial homology of the spot patterns provide a model to better understand the evolution of pattern development.

The remaining patterning elements also appear modular; however, they behave differently from the spot patterns with regards to gene expression correlation. Notably, we do not find any gene expression foreshadowing the *D. guttifera* wide shade on the abdomen; it may either be a gene that we did not test for, or there was no *in situ* hybridization signal because the expression levels were below the detection limit.

To fully understand the role of each gene in these three species’ color pattern development, we must utilize RNA interference and gene overexpression, as well as CRISPR approaches. Transgenic methods are established in *D. guttifera*; however, such procedures are not available for *D. palustris* and *D. subpalustris*. Pursuing the development of such approaches will facilitate a robust investigation of the mechanisms underlying these three species’ morphological diversity. Our understanding of color pattern development is far from complete; however, continuing to study these three fruit flies will help us connect the dots.

## Acknowledgments

We thank Dr. Rupali Datta for valuable comments on the manuscript.

